## supplement file S1 for "The theory of massively repeated evolution and full identifications of Cancer Driving Nucleotides (CDNs)"

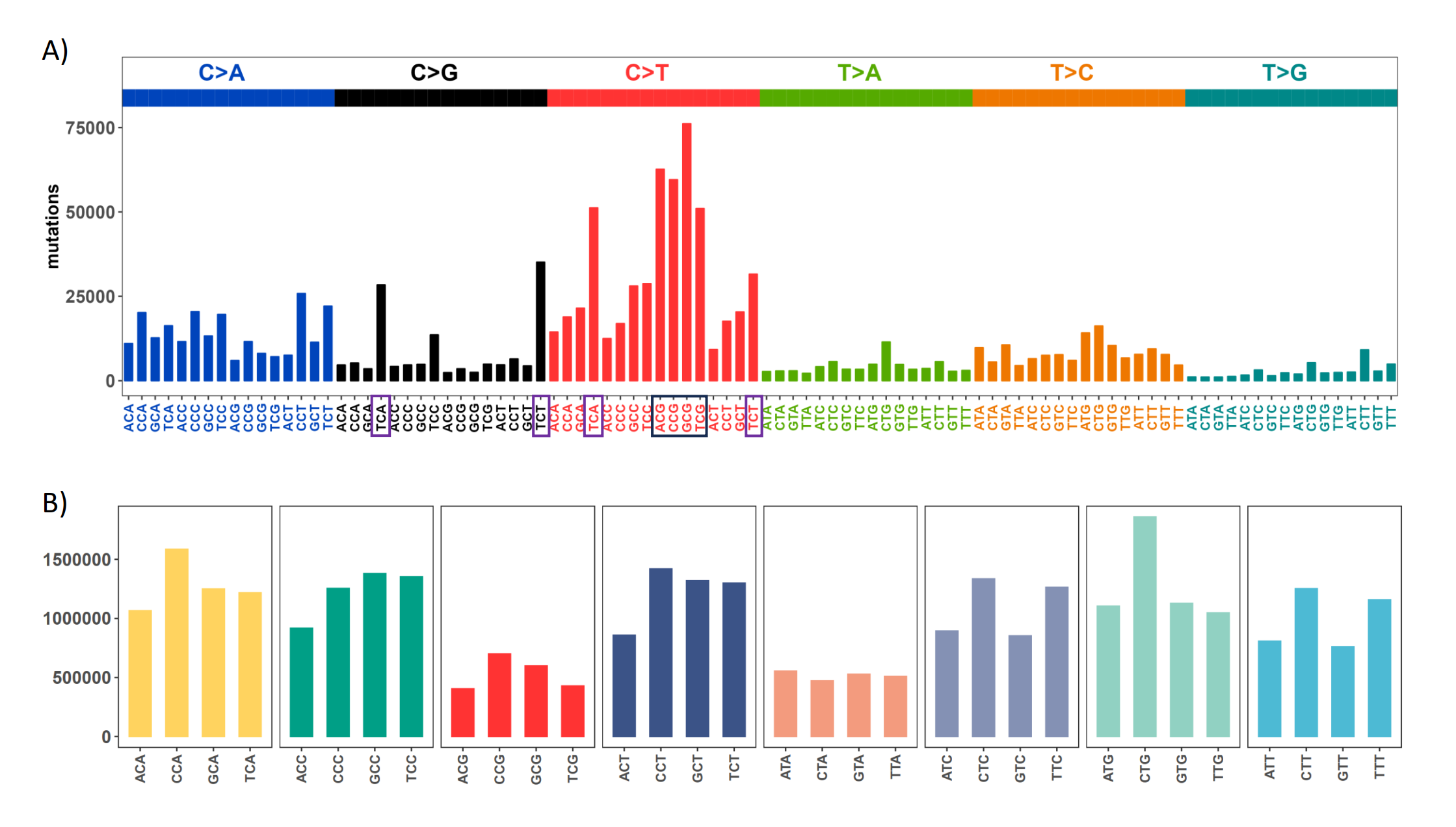

**Figure S1**: Mutation and context landscape across 12 cancer types.

**(A)** Single nucleotide changes within a local context of 3 bp across 12 cancer types. Mutations are grouped into 6 nucleotide change directions based on base complementarity, with colors representing each type. The Y-axis indicates the total number of each change type across the 12 cancers. The black box highlights the most abundant C>T (complementary G>A) changes occurring at CpG sites, while purple boxes designate the second-most prevalent nucleotide change at TCW (W = A or T) context which is potentially associated with APOBEC family of cytidine deaminases.

**(B)** Context abundance within the human reference genome. For each coding region site, we extract the local context by extending 1 bp to either side. Contexts are then collapsed into 32 categories with centered base being C or T based on base complementarity. The Y-axis shows the site count for each context along the X axis. The four red bars (ACG, CCG, GCG, TCG) highlight the abundance of CpG sites in human genome.

1. **Literature support for CDNs identified in breast cancer.**

Verification of site level positive selection in cancer genome has primarily focused on canonical cancer drivers. For non-canonical candidates with experimentally proven tumorigenic activity, CDN sites within these genes emerge as potential key drivers due to their statistically stronger selective advantage. Under this premise, we search for literature evidence for genes harboring CDN sites in breast cancer.

Among the 17 genes with CDN sites in breast cancer, 11 are recognized as canonical drivers by all three major driver gene lists. 4 genes (*CDC42BPA*, *ERBB3*, *KIF1B*, *NUP93*), despite lacking inclusion in canonical breast cancer driver lists, possess explicit experimental support indicating their driving roles in breast tumorigenesis. *HIST1H3B* has been recognized as a driver gene in breast cancer by IntOGen, corroborated by literatures supporting its association with breast cancer. The four mutation recurrences with R6C alteration in amino acid sequence in *RARS2* have been proposed to be linked to defects in mitochondrial transport, the explicit role of *RARS2* in breast cancer tumorigenesis remains to be explored.

**Table S1: literature support for CDN genes in breast cancer**

| **Gene Id** | **Gene Name** | **Support** |
| --- | --- | --- |
| ***AKT1*** | v-akt murine thymoma viral oncogene homolog 1 | ①②③ |
| ***CDC42BPA*** | CDC42 binding protein kinase alpha (DMPK-like) | (Unbekandt and Olson 2014; Collins et al. 2018; Kwa et al. 2021; Jiang et al. 2023) |
| ***CDH1*** | cadherin 1, type 1, E-cadherin (epithelial) | ①②③ |
| ***ERBB2*** | v-erb-b2 avian erythroblastic leukemia viral oncogene homolog 2 | ①②③ |
| ***ERBB3*** | v-erb-b2 avian erythroblastic leukemia viral oncogene homolog 3 | (Holbro et al. 2003; Xue et al. 2006; Hamburger 2008; Sithanandam and Anderson 2008; Stern 2008; Huang et al. 2010) |
| ***FGFR2*** | fibroblast growth factor receptor 2 | ①②③ |
| ***FOXA1*** | forkhead box A1 | ①②③ |
| ***GATA3*** | GATA binding protein 3 | ①②③ |
| ***HIST1H3B*** | histone cluster 1, H3b | ②(Xie et al. 2019; Wang et al. 2023)* |
| ***KIF1B*** | kinesin family member 1B | (Munirajan et al. 2008; Yu and Feng 2010; Liu et al. 2022) |
| ***KRAS*** | Kirsten rat sarcoma viral oncogene homolog | ①②③ |
| ***NUP93*** | nucleoporin 93kDa | (Bersini et al. 2020; Nataraj et al. 2022) |
| ***PIK3CA*** | phosphatidylinositol-4,5-bisphosphate 3-kinase, catalytic subunit alpha | ①②③ |
| ***PTEN*** | phosphatase and tensin homolog | ①②③ |
| ***RARS2*** | arginyl-tRNA synthetase 2, mitochondrial | (Wang et al. 2020)* |
| ***SF3B1*** | splicing factor 3b, subunit 1, 155kDa | ①②③ |
| ***TP53*** | tumor protein p53 | ①②③ |

Notes:

-The serial number corresponds to the inclusion of target gene in the following driver gene list:

① CGC Tier-1 list

② IntOGen

③ Bailey’s list

-The inclusion necessitates that the target gene is annotated as a cancer driver in breast cancer.

-*ambiguous, meaning the literature indicates an association between the candidate gene and breast cancer, but lacks explicit experimental evidence.

2. **All CDN sites with population allele frequency annotation.**

**Supplement file S2** contains five tables. “**CDN_sites.i>=20**” presents CDN sites with total hits ≥20 in each cancer type. “**CDN.Missense.thres_3**” provides all CDN sites analyzed in this study, ranked in decreasing order based on the highest recurrence across 12 cancer types. “**CDN.Missense.gnomAD**” presents the *gnomAD* population allele frequency of all missense CDNs. “**Synonymous_high_hits**” lists the synonymous mutations potentially under selection, while “**Synonymous_high_hits.gnomAD**” provides their corresponding allele frequency annotations from *gnomAD*.

3. **The impact of k for gamma-binomial model.**

From Eq. 2 in the main text, ***S_i_*** is affected by two terms: *G* and **nE(u)**, where *G* is:

$$G=L_{S}\cdot g\left( i,k \right)=L_{S}\cdot C_{k+i-1}^{i}\cdot\frac{1}{k^{i}}$$

***L_S_*** is a constant value given specific cancer type, here we demonstrate how $g\left( i,k \right)$ varies with respect to *i* and *k*, and elucidate why ***S_i_*** of *k* = 1 indicates the upper bound of CDN cutoff.

Considering the fold change of *G* from *i*-1 to *i*.

$$\varphi\left( i,k \right)=\frac{g\left( i,k \right)}{g\left( i-1,k \right)}=\frac{C_{k+i-1}^{i}\cdot\frac{1}{k^{i}}}{C_{k+i-2}^{i}\cdot\frac{1}{k^{i-1}}}$$

Eq. S8

**Fig. S2** illustrates how $\varphi\left( i,k \right)$ changes with *i* and *k*. With *k* range from 0.1 to 10, the curve of $\varphi\left( i,k \right)$ elucidates the extent to which *G* would impact ***S_i_*** with each increment of *i*. As detailed in **Section 4**, *k* will be >1 for biological significance. In such cases, $\varphi\left( i,k \right)$ will always be <1, meaning *G* will synergistically collaborate with **nE(u)** to decrease ***S_i_***. The diminishing impact of $\varphi\left( i,k \right)$ intensifies as *k* increases. A higher *k* value would suggest that, for most sites across the genome, mutability falls within a narrow range of >0. In an extreme case, when *k* = 10, $\varphi\left( i,k \right)$ = 0.145 at *i* = 20, which is 2 orders weaker than **nE(u)** for cancer types in TCGA. In practice, *k* is usually estimated to be between 2-5, depending on the cancer types being investigated (Table S2). Consequently, the reduction of ***S_i_*** with each increase of *i* is predominantly governed by **nE(u)**, and the ***S_i_*** values with *k* = 1 represent the upper limit driven solely by mutational force.

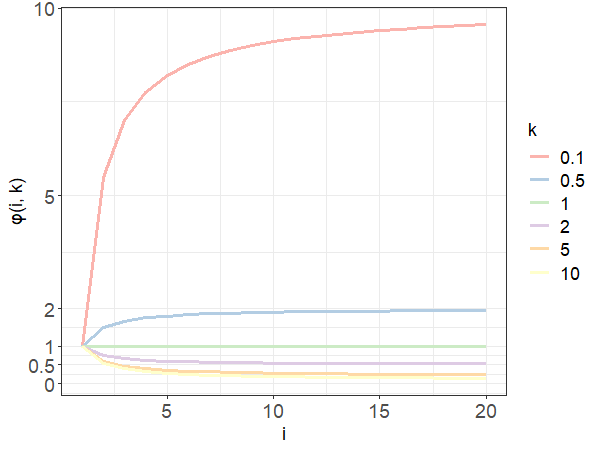

**Figure S2**: the trend of $\varphi\left( i,k \right)$ with each increase of recurrence (*i*, the x-axis) under different shape parameters of the gamma distribution (*k*, designated by different colors).

**Table S2**: *k* estimated from 12 cancer types.

| **Cancer type** | ***k*** |
| --- | --- |
| Breast | 5.05 |
| CNS | 2.59 |
| Endometrium | 5.49 |
| Kidney | 7.70 |
| Large intestine | 4.76 |
| Liver | 5.23 |
| Lung | 2.62 |
| Ovary | 4.30 |
| Prostate | 3.60 |
| Stomach | 4.17 |
| Upper-AD tract | 4.14 |
| Urinary tract | 6.14 |
| **merged set^*^** | **3.27** |

**Note:**

- Estimation of *k* is derived from negative binomial regression, based on synonymous changes aggregated by the 3 bp local context at mutated sites across all coding genes. The estimation method is implemented in package dndscv.

-* The merged set contains mutation information from all 12 cancer types.

4. **The impact of negative selection on shape parameter *k*.**

Across various studies aiming to depict mutability variation across genome under the gamma distribution, the shape parameter *k* is always pivotal. Adopting a dichotomous perspective, we inquire into how *k* compares to 1 under large sample sizes (**n** ≥ 10^6^). This inquiry is fundamentally linked to the prevalence of negative selection across the cancer genome, as the observed mutation abundance is an amalgamation of mutational and selection forces. In scenarios where purifying selection extensively operates throughout the genome in cancer evolution, even for synonymous sites (Sharp and Li 1987; Plotkin and Kudla 2011; Gartner et al. 2013; Chu and Wei 2019), most genomic sites would not exhibit mutations, resulting in *k* being ≤1. In attempts to detect negative selection signals in cancer, researchers typically identify only a limited number of genes (Luo et al. 2008; Eynden et al. 2016; Zapata et al. 2018; Bányai et al. 2021). In a CRISPR-Cas9 loss-of-function screen covering 16,540 genes conducted across 558 cancer cell lines, only approximately 6% of genes are under strong negative selection in at least 90% of the cell lines (De Kegel and Ryan 2019). In an in-house mutation accumulation experiment carried out in HCT116 (a human colorectal carcinoma cell line, data not published), the proportion of mutations under strong negative selection is 0.66% with a selection coefficient (*s*) of -0.6 (indicating that the survivability of the mutant is 40% of the wildtype). These evidences, in concordance with quasi-neutrality of cancer evolution, suggest that purifying selection is indeed rare in cancer evolution. The mutability for the majority of genomic sites is greater than 0, with shape parameter *k* > 1.

5. **Detailed derivation for negative binomial distribution and approximation of *i^*^*/n.**

The derivation of Eq. 4 from joint distribution of Gamma-Poisson distribution is well presented in statistically analysis. Here, we assume that the mutation recurrence (*i*) observed at site level across the genome follows a Poisson distribution of $Pois\left( i|\lambda\right)$, where the expected number of mutation recurrence $\lambda$ follows a Gamma distribution of $Gamma\left( \lambda|k,\theta\right)$, with *k* and $\theta$ being the shape and scale parameters, respectively. Then, the joint probability density function for *i* can be expressed as:

$$f\left( i|k,\theta\right)=\int Pois\left( i|\lambda\right)\cdot Gamma\left( \lambda|k,\theta\right)\cdot d\lambda$$

$$=\int\frac{\lambda^{i}e^{-\lambda}}{i!}\cdot\frac{1}{\Gamma\left( k \right)\theta^{k}}\lambda^{k-1}e^{-\frac{\lambda}{\theta}}\cdot d\lambda$$

$$=\frac{\theta^{-k}}{i!\Gamma\left( k \right)}\int\lambda^{i}e^{-\lambda}\cdot\lambda^{k-1}e^{-\frac{\lambda}{\theta}}\cdot d\lambda$$

$$=\frac{\theta^{-k}}{i!\Gamma\left( k \right)}\int\lambda^{\left( i+k \right)-1}e^{-\left( 1+\frac{1}{\theta} \right)\lambda}\cdot d\lambda$$

Eq. S9

Now, we make use of the probability density function of Gamma distribution,

$$\int Gamma(\lambda|k,\theta)=1$$

Which is:

$$\int\frac{1}{\Gamma\left( k \right)\theta^{k}}\lambda^{k-1}e^{-\frac{\lambda}{\theta}}\cdot d\lambda=1$$

Therefore,

$$\int\lambda^{k-1}e^{-\frac{\lambda}{\theta}}\cdot d\lambda=\Gamma\left( k \right)\left( \frac{1}{\theta} \right)^{-k}$$

Eq. S10

Comparing with Eq. S10, Eq. S9 can be rewrite as:

$$f\left( i|k,\theta\right)=\frac{\theta^{-k}}{i!\Gamma\left( k \right)}\cdot\left( 1+\frac{1}{\theta} \right)^{-\left( i+k \right)}\Gamma\left( i+k \right)$$

$$=\frac{\theta^{-k}}{i!\Gamma\left( k \right)}\cdot\left( \frac{\theta}{1+\theta} \right)^{i+k}\Gamma\left( i+k \right)$$

$$=\frac{\Gamma\left( i+k \right)}{\Gamma\left( i+1 \right)\Gamma\left( k \right)}\left( \frac{1}{1+\theta} \right)^{k}\left( \frac{\theta}{1+\theta} \right)^{i}$$

Eq. S11

Note that the mean for Gamma distribution is $k\theta=nE\left( u \right)$, which leads to $\theta=\frac{nE\left( u \right)}{k}$. Then, the negative-binomial form of Eq. S11 could be further expressed as:

$$f\left( i|k,\theta\right)=\frac{\Gamma\left( i+k \right)}{\Gamma\left( i+1 \right)\Gamma\left( k \right)}\left( \frac{k}{k+nE\left( u \right)} \right)^{k}\left( \frac{nE\left( u \right)}{k+nE\left( u \right)} \right)^{i}$$

$$=\frac{\Gamma\left( i+k \right)}{\Gamma\left( i+1 \right)\Gamma\left( k \right)}k^{k}\left[ nE\left( u \right) \right]^{i}\left[ k+nE\left( u \right) \right]^{-k-i}$$

Which is Eq. 6 from the main text.

With *k* = 1, $nE\left( u \right)=k\theta=\theta$, Eq. S11 then transforms to:

$$f\left( i|1,\theta\right)=\frac{\Gamma\left( i+1 \right)}{\Gamma\left( i+1 \right)\Gamma\left( 1 \right)}\left( \frac{1}{1+\theta} \right)^{1}\left( \frac{\theta}{1+\theta} \right)^{i}$$

$$=\left( \frac{1}{1+\theta} \right)\cdot\left( 1-\frac{1}{1+\theta} \right)^{i}$$

$$=\left( \frac{1}{1+nE\left( u \right)} \right)\cdot\left( 1-\frac{1}{1+nE\left( u \right)} \right)^{i}$$

Which is a geometric distribution with $p=\frac{1}{1+nE\left( u \right)}$.

For the approximation of *i^*^*/**n**, we let *ε* = 1, then Eq. 10 from main text could be rewritten as:

$$i^{*}\cdot\log\left( \frac{1}{1+\frac{1}{nE\left( u \right)}} \right)=\log\left( \frac{1}{L_{A}} \right)$$

$$i^{*}\cdot\log\left( 1+\frac{1}{nE\left( u \right)} \right)=\log\left( L_{A} \right)$$

Eq. S12

For the left side of Eq. S12, we use the first-order Tayler expansion,

$$i^{*}\cdot\log\left( 1+\frac{1}{nE\left( u \right)} \right)\sim i^{*}\cdot\frac{1}{nE\left( u \right)}$$

Substitute this to Eq. S12, we have:

$$\frac{i^{*}}{n}=\log(L_{A})\cdot E\left( u \right)$$

Which is Eq. 11 from main text.

6. **Probing mutation rate variation with large samples.**

With large sample size sequenced, the data will yield an additional benefit by revealing the evolution of the mutation rate itself. Given that the mutation rate per site is extremely small, the evolution of mutation rate itself has been a most challenging issue (André and Godelle 2006; Lynch 2010; Lynch 2011; Lynch et al. 2016; Ruan et al. 2020; Wei et al. 2022). In particular, without the check of selection, the mutation rate is liable to be trapped in the runaway evolution (Ruan et al. 2020).

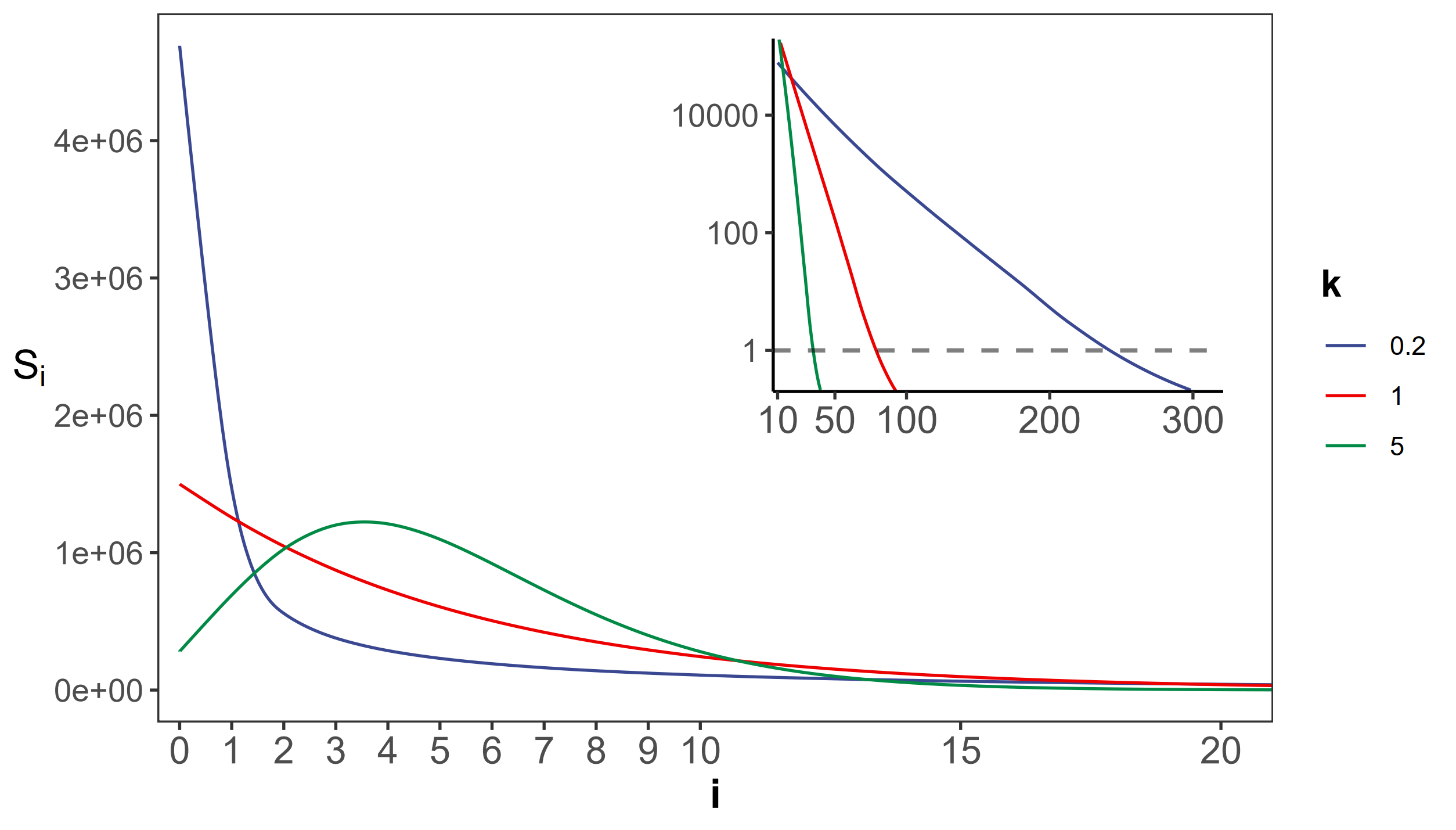

**Figure S4:** the gamma distribution of recurrences (*i*) under different shapes. With **E(u)** = 5×10^-6^, we set the shape parameter *k* to 0.2, 1 and 5, represented by three distinct colors. The site number of synonymous recurrence *i* (***S_i_***) is indicated on Y-axis. In the context of a large sample size (**n**=10^6^), the ***S_i_*** distribution clearly distinguishes between different *k* values, mitigating the overdispersion issue encountered in smaller sample sizes. The inset depicts the distribution on a log10 scale for *i* ≥ 10, with a horizontal dashed line indicating $\boldsymbol{S}_{\boldsymbol{i}^{\boldsymbol{*}}}=1$, where *i^*^* is the CDN cutoff.

The theory of mutation rate of evolution should be based on the distribution of the per-site mutation rate across the genome. However, the empirical data so far only yield the mean. In particular, the spectrum of ***S_i_***’s for *i*’s close to 1 would be most informative about the evolution of the mutation mechanism. **Fig. S4** shows the ***S_i_*** spectrum with *k* = 0.2, 1 or 5 in a Gamma distribution. Note the mode of the distribution (i.e., the peak of the curve) among the 3 curves, which is at 0 or >0 depending on whether *k* ≤ 1 or > 1. Clearly, the observed ***S_i_***’s can distinguish among the three distributions only when **n** is very large. The implications of such distributions for the theory of mutation rate evolution are addressed in Discussion.

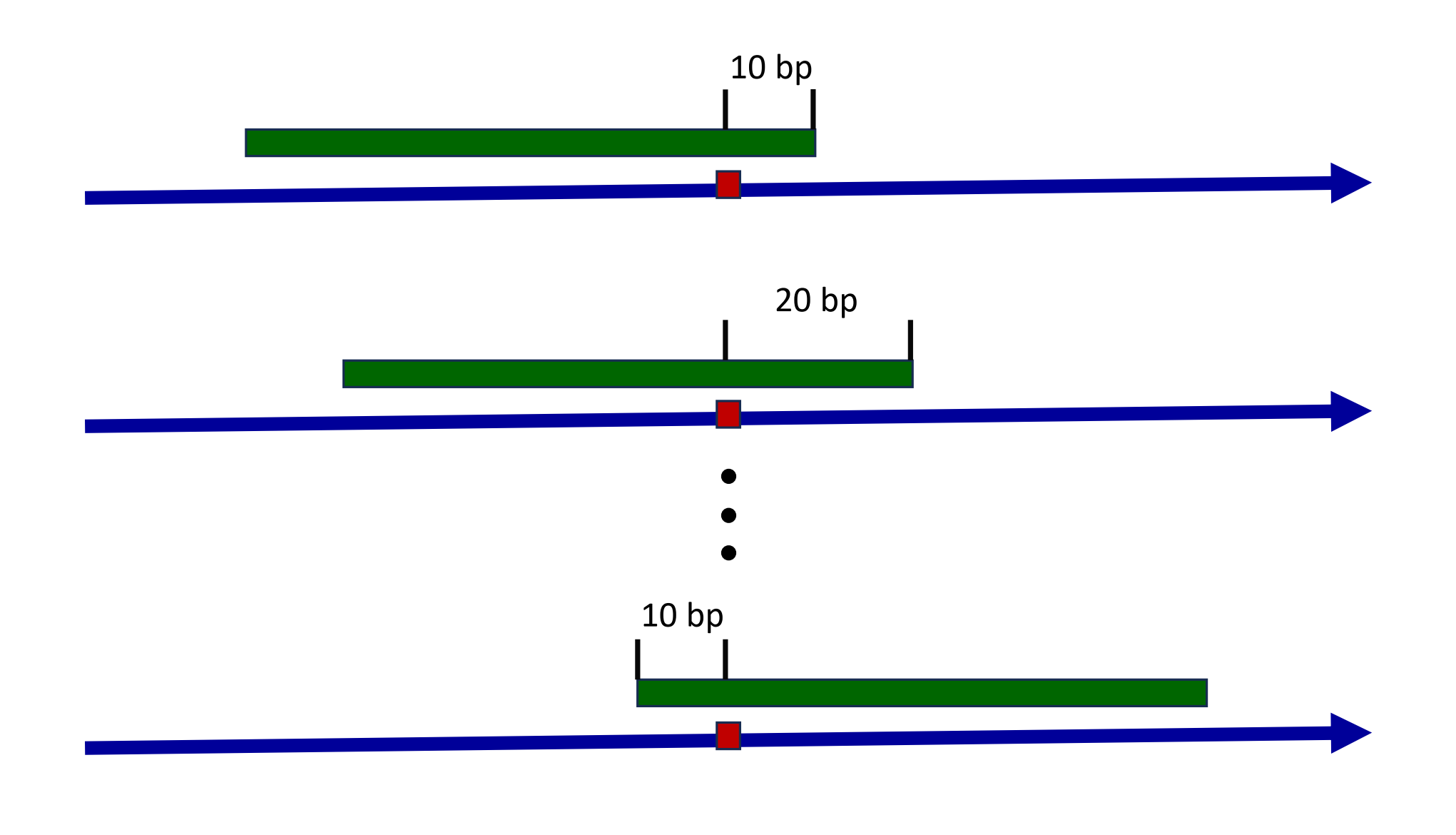

**Figure S5:** Sliding window to explore the consensus sequences between recurrence sites. The blue arrow indicates the positive strand of reference genome, with a mutated site highlighted by the red box. The green strip represents a sliding window covering the mutated site. With each stride of 10 bp, we extract the sequence context and conduct a pairwise comparison between all recurrence sites. Presence of consensus motifs would skew the Hamming distance distribution away from the expected Poisson model, reflecting non-randomness.
